# Chromatin profiling for everyone: FFPE-CUTAC for the theory and practice of modern molecular biology

**DOI:** 10.64898/2025.12.24.696395

**Authors:** Yiling Xu, Steven Henikoff, Kami Ahmad

**Affiliations:** Fred Hutchinson Cancer Center, Seattle WA; Howard Hughes Medical Institute, Chevy Chase MD

## Abstract

In 2025, together with the Fred Hutch Summer Undergraduate Research and Summer High School Internship programs, we developed and implemented a laboratory genomics research experience to introduce students to modern molecular biology techniques and bioinformatics. The course centered around using a new method we had developed in 2023 that uses readily available fixed tissue sections on glass slides. Students performed a series of steps to tagment genomic locations of RNA Polymerase II and then used PCR to enrich libraries for next-generation sequencing in a core facility. Students then visualized their data in genomic browser tracks and assessed the results. At the end of the summer, students prepared and presented their work and experiences in seminar format to their cohorts. Overall, the technical simplicity of on-slide chromatin profiling introduced the students to laboratory practice and current techniques in genomics, bioinformatics, and medical sciences.

## Introduction

Standard clinical practice for collecting tissue specimens begins with formalin fixation and paraffin embedding (FFPE), then sectioning the paraffin block on a microtome, followed by transfer of thin sections to microscope slides for staining and examination by a pathologist. The remaining FFPE blocks have been accumulating in hospitals and clinics world-wide by the billions, resulting in valuable resources that are often annotated with anonymized patient metadata that can be used for DNA, RNA or protein molecular analyses^1^. We have previously introduced a variation of our CUT&Tag chromatin profiling method, Cleavage Under Targets and Tagmentation (CUTAC), that is especially well-suited for FFPEs, in which the targets are gene regulatory elements occupied by RNA Polymerase II (Pol II)^2-5^. Pol II produces all mRNAs and most other types of RNA in an organism, and so FFPE-CUTAC maps the basic machinery that drives gene expression making it well-suited for active teaching. While high school and undergraduate students are introduced to the Central Dogma of molecular biology (DNA is transcribed to make mRNA which is translated to make proteins), FFPE-CUTAC puts this into laboratory practice and connects it to current practices in genomics, bioinformatics, and medical sciences.

During the summer of 2025 we hosted 2 high-schoolers and 2 sophomore undergraduates through the Fred Hutch Summer internship programs. Students were selected and paid to spend about 6 weeks embedded in research labs at the Hutch. Our four summer students were tasked with performing FFPE-CUTAC each week under the supervision of a lab mentor, and these experiments were used to discuss principles and techniques during incubation steps. Each student prepared and presented their work and experiences in seminar format to the entire internship cohort and their mentors at the end of the summer. Overall, the technical simplicity of FFPE-CUTAC enabled these four students to produce research-quality data. We plan to develop a module scaled for a high school curriculum for the 2026-2027 school year that can be broadly implemented.

## Methods

We made two changes to the FFPE-CUTAC protocol (https://www.protocols.io/view/cutac-for-ffpes-14egn292zg5d/v4) for new students. First, we composed a checklist of steps that students could follow during their lab time. This helped students track their work through the multi-day protocol. Second, Step 19 of the protocol uses a buffer containing N,N-dimethylformamide (DMF) for tagmentation; Environmental Health and Safety regulations do not permit high-schoolers to use this potentially hazardous chemical. However, our earlier work showed that the enhanced tagmentation in CUTAC protocols is primarily due to the low ionic strength of this buffer^2^. Therefore, the two undergraduate students performed FFPE-CUTAC with DMF-free buffer and obtained indistinguishable results from using buffer with DMF. All further experiments in the summer for the four students used DMF-free buffer. This option is provided in the protocol.

## Recipes

*Cross-link reversal buffer (100 mL) 800 mM Tris-HCl, 0*.*2 mM EDTA pH8*.*0:*

80mL 1M Tris-HCl pH 8.0

20mL dH_2_O

40µl 0.5 mM EDTA

*Triton-Wash buffer (50 mL) 20 mM HEPES pH 7*.*5, 150 mM NaCl, 0*.*5 mM spermidine, 0*.*05% Triton-X100* + *protease inhibitor tablet:*

1 mL 1 M HEPES pH 7.5

1.5 mL 5 M NaCl

250 µl 10% Triton-X100

12.5 μl 2 M spermidine

20 µl 0.5M EDTA

Bring the final volume to 50 mL with dH_2_O

Add 1 Roche Complete Protease Inhibitor EDTA-Free tablet

Store the buffer at 4°C for up to 2 days.

*Primary antibody solution (100 µl for every 2-4 samples):*

4 µl RNA Polymerase II-Ser5p: (Cell Signaling Technologies (D9N5I) mAb #13523)

96 µl Triton-Wash buffer (1:25)

*Secondary antibody solution (100 µl for every 2-4 samples):*

4 µl guinea pig anti-rabbit (Antibodies Online)

96 µl Triton-Wash buffer (1:25)

*Protein AG-Tn5 solution (100 µl for every 2-4 samples):*

5 µl Protein AG-Tn5 (Epicypher cat. no. 15-1117)

95 µl Triton-Wash buffer (1:20)

*CUTAC-DMF Tagmentation buffer (50 mL) 10 mM TAPS, 5 mM MgCl*_*2*_, *20% DMF:*

39.3 mL dH_2_O

10 mL N,N-dimethylformamide

0.5 mL 1 M TAPS pH 8.5

250 µl 1 M MgCl_2_

Store the buffer at 4°C for up to 1 week.

*CUTAC-noDMF Tagmentation buffer (50 mL) 10 mM TAPS, 10 mM MgCl*_*2*_:

49.3 mL dH2O

0.5 mL 1 M TAPS pH 8.5 250 µl 1 M MgCl_2_

Store the buffer at 4°C for up to 1 week.

*10mM TAPS (100 mL):*

1mL 1M TAPS pH8.5

99mL dH_2_O

*TAPS wash buffer (100 mL) 10 mM TAPS, 0*.*2 mM EDTA:*

99 mL dH2O

1mL 1 M TAPS pH 8.5

40 µl 0.5 M EDTA

*1% SDS/ProtK Release solution (100 µl for 16 samples):*

10 µl 10% SDS

1 µl 1 M TAPS pH 8.5

79 µl dH_2_O

Just before use add 10 µl Thermolabile Proteinase K (NEB cat. no. P8111S).

*5% Triton Mix (1 mL):*

500 µl 10% Triton-X100

500 µl dH_2_O

Store at room temperature

*Barcoded i5 primers (10 µM solution):*

**5’-AATGATACGGCGACCACCGAGATCTACAC(8-bp i5 barcode)TCGTCGGCAGCGTCAGATGTGTAT-3’**

*Barcoded i7 primers (10 µM solution):*

**5’-CAAGCAGAAGACGGCATACGAGAT(8-bp i7 barcode)GTCTCGTGGGCTCGGAGATGTG-3’**

## Materials

Plastic film rectangles (Cut from food-service plastic wrap, *e*.*g*. Reynolds 912 Razor blades

Forceps

FFPE mouse brain tumor sample slides each with 4-5 tissue sections

### Protocol

*Day 1*

1) Deparaffinization (< 2 hours):

Place slides in a Coplin jar containing Safe Clear II and hold for 10 min at room temperature. Transfer slides to a Coplin jar containing 100% ethanol and hold for 3-5 min.

Transfer slides to a Coplin jar containing 50% ethanol and hold for 3-5 min. Transfer slides to a Coplin jar containing 10% ethanol and hold for 3-5 min.

Transfer slides to a plastic Coplin jar containing cross-link reversal buffer and incubate at 85°C for 1 hr.

Place the plastic Coplin jar into ice-cold water to cool down.

2) Antibody binding (∼20 min prep + incubation overnight):

Take slides out one by one. For each slide, add 25-50 µl primary antibody solution. Cover the clear portion of the slide with a rectangle of plastic film using surface tension to spread the liquid. Place vertically side-by-side in a 30-slide holder moist chamber. Incubate at 4°C O/N.

*Day 2*

3) Bind secondary antibody (∼30 min prep + 1.5 hr incubation):

Place on a wet paper towel pad to wash the back of the slide, then gently rinse the top 1-2 times with 1 ml Triton-Wash buffer. For each slide add 25-50 µL 1:25 secondary antibody solution. Cover with plastic wrap rectangles and place in 30-slide holders in moist chamber. Incubate at RT for at least 1 hr.

4) Bind pAG-Tn5 (∼30 min prep + 1.5 hr incubation + 20min):

Place on a wet pad to wash the back of the slide and then gently rinse the top with 1 ml Triton-Wash buffer. For each slide add 25-50 µL pAG-Tn5 solution. Cover with plastic wrap squares and place in 30-slide holders in a moist chamber. Incubate at RT for at least 1 hr. Remove film and gently rinse the top 1-2 times with a 1 mL Triton-Wash-EDTA. Drain well and transfer to Coplin jars containing 10 mM TAPS pH 8.5 for 10 min.

5) Tagmentation (1 hr incubation + 20 min):

Drain and wick off excess liquid. Transfer slides from 10 mM TAPS into plastic Coplin jars with cold tagmentation buffer. Incubate 55°C 1 hr in a water bath.

Place the Coplin jars in ice water to cool down. Transfer the slides into Coplin jars containing 10 mM TAPS pH8.5, 0.2 mM EDTA (TAPS Wash) to hold at 4°C O/N.

*Day 3*

6) Fragment release (30 min prep + 1.5 hr incubation):

Just before use mix 1.1% SDS 10 mM TAPS with Thermolabile Proteinase K (1:10). Drain and blot off excess using a Kimwipe.

First add 7 µL ProtK/SDS to each tube. Add 3 µL ProtK/SDS per sample to the tissue using a pipette tip and scrape everything into a pile using a razor blade. Then transfer everything to the tube with #5 forceps. There should be ∼10 µl ProtK/SDS total in each tube.

Incubate 37°C 30 min -> 58°C 30 min.

While tubes are still on the cycler add 30 µL 5% Triton-X. Incubate 37°C 30 min.

At this point, the total volume of each sample is ∼40 µl (10 µl + 30 µl). For smaller size tissue, use 5 µl ProtK/SDS + 15 µl 5% Triton-X. Adjust in proportion, depending on tissue size.

7) PCR (50 min):

Add 2 µl each barcoded i5 and i7 primers, using a different i5 or i7 barcode for each sample to be pooled together for sequencing.

Chill on ice. Add 25 µl NEBNext (New England Biolabs cat. no. E7400L). Use PCR CNT-EXT (30 sec annealing and 1m extension) for 13 total cycles. Cycle 1: 58°C for 5 min (gap filling)

Cycle 2: 72°C for 5 min (gap filling)

Cycle 3: 98°C for 5 min

Cycle 4: 98°C for 10 sec

Cycle 5: 63°C for 30 sec

Cycle 6: 72°C for 1 min

Repeat Cycles 4-6 12 times

Hold at 8°C

To avoid possible inhibition of the PCR, we used only a fraction of each sample or section. Sample 1-8: 2 µl sample + 18 µl water (2 µl out of 40 µl = 5% of the total tissue on the slide) Sample 9-16: 4 µl sample + 16 µl water (4 µl out of 40 µl = 10% of the total tissue on the slide) Save the rest for future use (reruns or replicates). Store at 4°C.

8) SPRI cleanup:

Add 65 µl SPRI beads. Wash twice with 200 µl 80% ethanol. Quick spin and remove the last drop. To avoid drying, immediately elute in 22 µl 0.1xTE8. Use 2 µl for Tapestation analysis, and the remaining 20 µl for sequencing.

## Results

In previous versions of this protocol we obtained high-quality results mapping Pol II in FFPE preparations of human tissue^5^ and of mouse brains^4^ (**Figure 1**). We have streamlined and optimized this protocol, which we describe here. These experiments were performed as part of a wet-lab education experience for summer students. We obtained slides of brain sections from mice injected with constructs for YAP1 fusion gene expression^6^ and performed FFPE-CUTAC on those slides. Each slide had 4-5 sections of tissue that were incubated together through CUTAC steps, and after tagmentation each section was scraped and transferred to a tube DNA release and barcoded PCR. Library enrichment was first assayed by capillary gel electrophoresis on an Agilent Tapestation, and successful reactions produced a 200 bp band that corresponds to an ∼50 bp insert with 138 bp of adapter sequences (**Figure 2**). Occasional reactions showed no measurable library band, however if parallel reactions were successful based on the gel read-out, often the blank reactions also yielded successful libraries. In these cases, all samples were submitted for DNA sequencing. Barcoded samples were pooled based on measured library concentrations with the intention of producing ∼10 million fragments per sample and sequenced on an Illumina NovaSeq X machine in dual-index paired-end mode. Reads were mapped to the mouse Mm10 genome assembly. Table 1 shows sequencing statistics for replicate samples, including one (P03) that showed no detectable library after enrichment, but nevertheless produced 13 million reads in the pool. Browser tracks of these four samples showed peaks of reads at the highly expressed histone genes (**Figure 3**).

**Figure 1.**
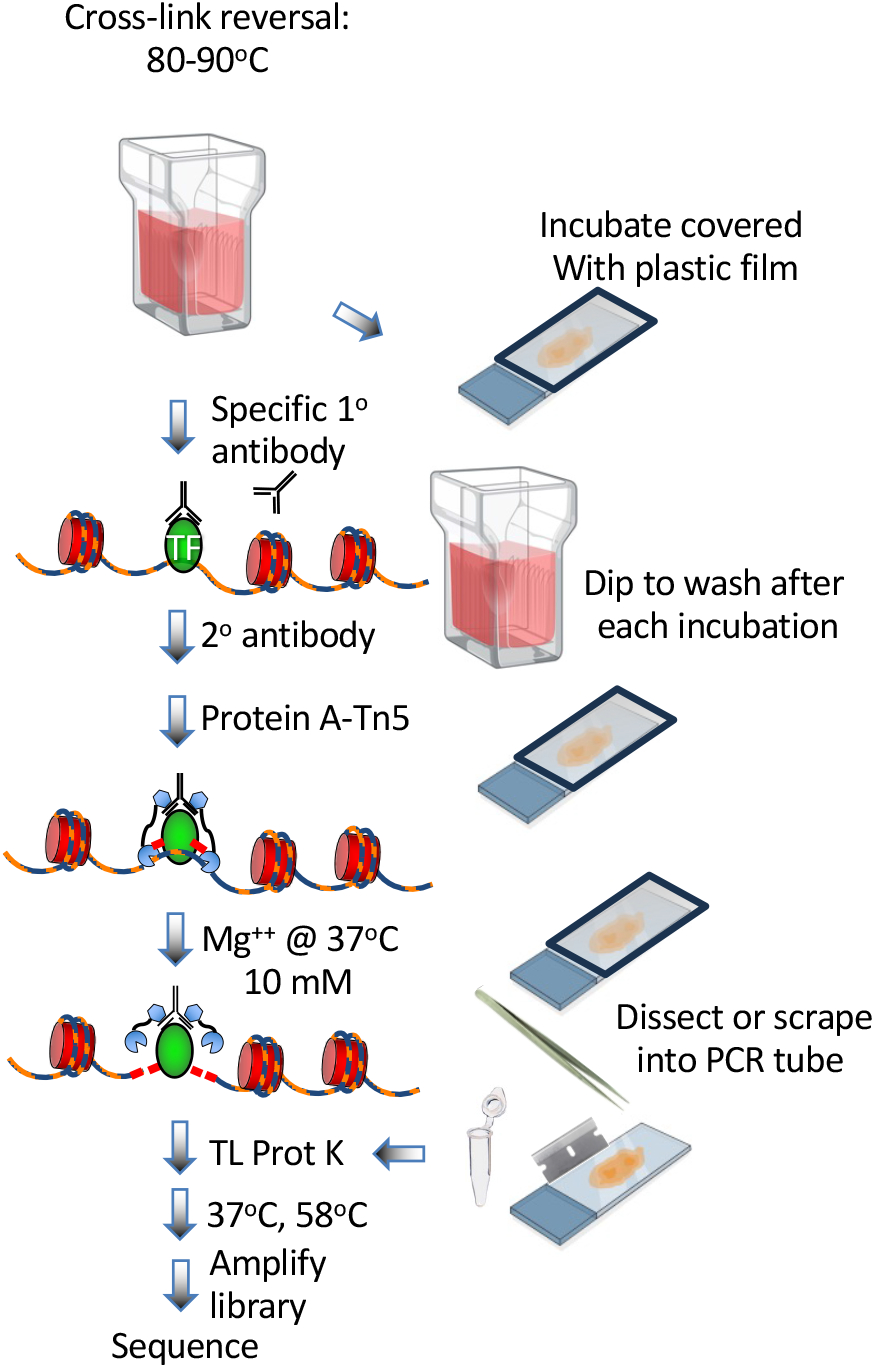
Overview of the FFPE-CUTAC protocol.

**Table 1.** Representative sequencing statistics.

| Sample | Raw_reads | Clipped_reads | mm10_fragments | ecoli_fragments | mm10_fraction_mapped | chrM_fraction | ecoli fraction | DUPs | ELS |
| --- | --- | --- | --- | --- | --- | --- | --- | --- | --- |
| 1 | 19,579,836 | 19,506,366 | 17,554,458 | 31,215 | 0.900 | 0.047 | 0.0016 | 0.937 | 1,103,963 |
| 2 | 17,246,414 | 17,177,260 | 15,366,973 | 30,904 | 0.895 | 0.027 | 0.0018 | 0.934 | 1,012,985 |
| 3 | 13,466,026 | 13,426,404 | 12,366,558 | 19,646 | 0.921 | 0.043 | 0.0015 | 0.926 | 913,866 |
| 4 | 10,026,912 | 9,993,769 | 8,474,775 | 19,528 | 0.848 | 0.047 | 0.0019 | 0.884 | 986,151 |

DUPs – duplicate reads, inferred by MarkDuplicates (Picard) package
(https://broadinstitute.github.io/picard/)
ELS – estimated library size, calculated by MarkDuplicates (Picard)

**Figure 2.**
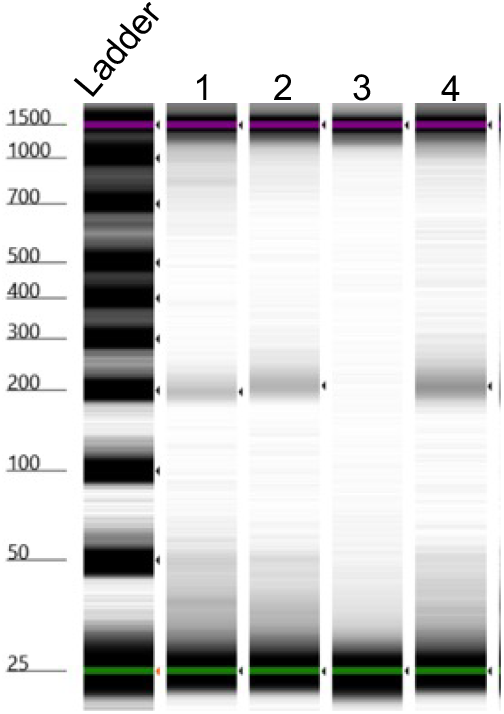
Gel analysis of sequencing libraries generated by FFPE-CUTAC. Each PCR reaction was run on an Agilent Tapestation 4200 and quantified for Illumina sequencing. Fragment sizes are estimated by the electronic ladder shown on the left. Productive experiments typically produce a product ∼200 bp in size. Although occasional reactions have low yields, these often produce enough sequencing for efficient genome mapping.

**Figure 3.**
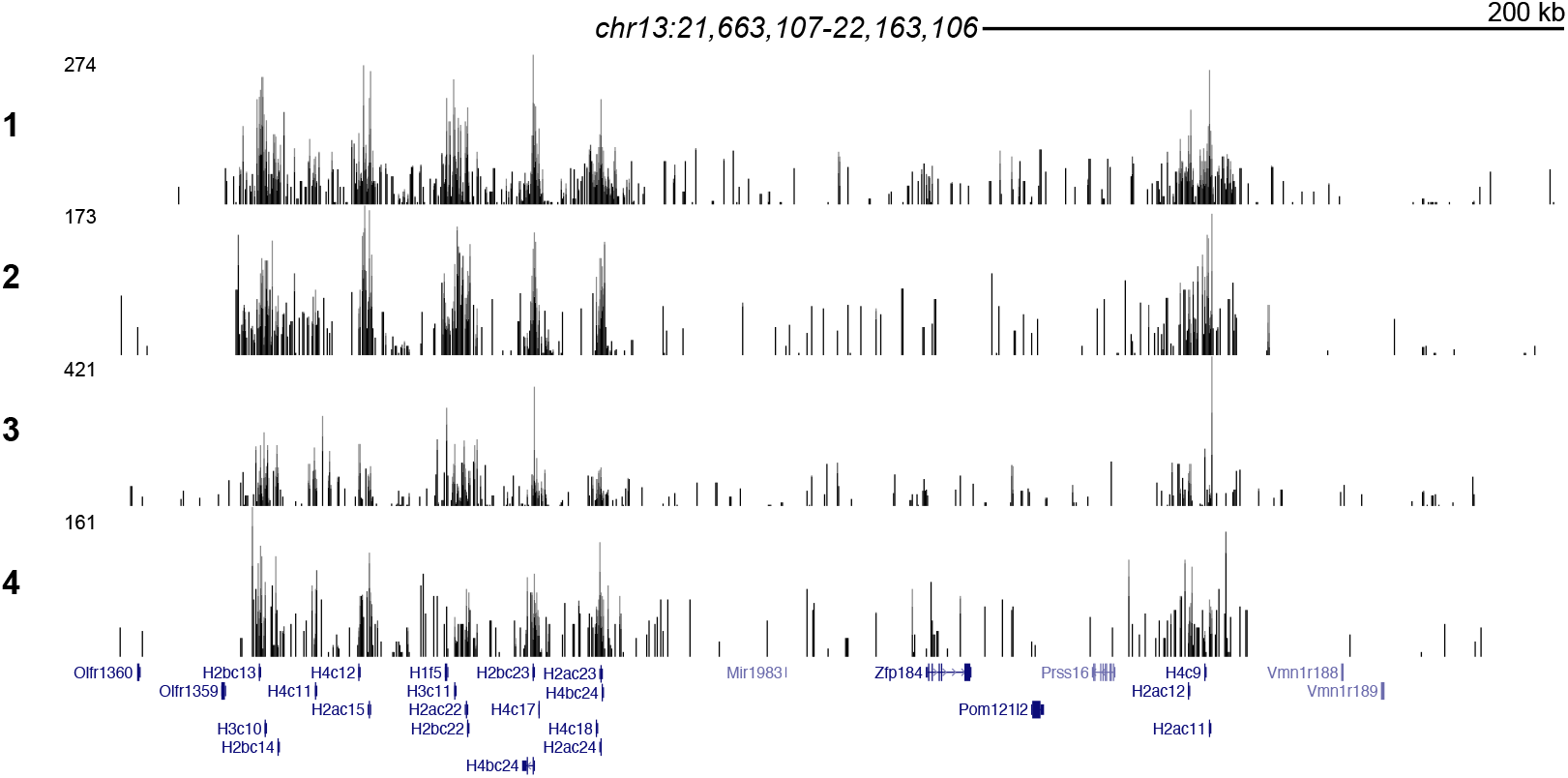
RNAPII enrichment at histone genes in the mouse genome. Mapped fragments from the four replicate libraries (Figure 2) are shown across a region of the mouse genome containing multiple highly expressed histone genes. RNA Polymerase II signal forms peaks at the promoters of the active genes.

## Discussion

Here we have described a chromatin profiling protocol based on a method that we routinely perform in the laboratory using standard pathology samples, but customized for the classroom and implemented during a 6-week summer course for undergraduate and high-school students. The intent of this article is both to disseminate our customized protocol to educators and to provide an example of a detailed protocol article describing an extension of a published protocol targeting a specific community. Our protocol is linked to “CUTAC for FFPEs”, which was first posted online in 2023 and currently on Version 5 (Ref. 7), thus offering the opportunity for teachers and professors who adopt this protocol to easily update their curriculum in future years. There has been an unmet need for a publication venue that can be used to extend, update, or customize published experimental or computational methods that are useful but technologically incremental. We hope that our example will encourage the dissemination of practical methods and step-by-step protocols that otherwise would not be adopted beyond the laboratory that developed them.

## Appendix 1

**Checklist for FFPE-CUTAC experiments**

*Based on https://www.protocols.io/view/cutac-for-ffpes-14egn292zg5d/v4*

1. **Deparaffinization**
  – Safeclear 2 in Coplin jar
  – EtOH series (100%, 50%, 10%) in Coplin jars
  – Xlink reversal in Coplin jar (85° 1hr)
2. **1° antibody binding**
  – 25 µL with saran wrap cover
3. **2° antibody binding**
  – rinse
  – 25 µL with saran wrap cover
4. **pAGTn5 binding**
  – inse
  – 25 µL pAGTn5 dilution
  – Coplin jar to rinse
5. **Tagmentation**
  – in Coplin jar (55° 1hr)
6. **Fragment release**
  – scrape into tubes
  – SDS/protease K incubation in PCR machine
  – quench with triton-X100
7. **PCR**
  – set up reactions on ice
  – run PCR program
8. **Ampure (SPRI) cleanup**
  – add SPRI beads
  – use magnets, wash with 80% EtOH
  – elute with 10 mM Tris pH8
9. **tapestation**
  – run!

## Appendix 2

**Molecular biology topics for background and discussion**

How big is the human genome? What’s in it? (a very small fraction is genes)

Central Dogma of Molecular Biology from the perspective of machinery (What is DNA polymerase? What is RNA polymerase? What is the ribosome?)

Where do the enzymes of molecular biology come from (PCR enzymes from thermophiles, transposases from genetic parasites)

*An aside: most of genomes are parasitic DNAs*

Enzymology of transposons Enzymology of PCR

How does Illumina sequencing work? (What is a flowcell?; What is chemical sequencing?)

How do we map sequence to genome assemblies? (What are the concepts of matching strings, with a scoring matrix)

